# Reprogramming, oscillations and transdifferentiation in epigenetic landscapes

**DOI:** 10.1101/193599

**Authors:** Bivash Kaity, Ratan Sarkar, Buddhapriya Chakrabarti, Mithun K. Mitra

## Abstract

Waddington’s epigenetic landscape provides a phenomenological understanding of the cell differentiation pathways from the pluripotent to mature lineage-committed cell lines. In light of recent successes in the reverse programming process there has been significant interest in quantifying the underlying landscape picture through the mathematics of gene regulatory networks. We investigate the role of time delays arising from multistep chemical reactions and epigenetic rearrangement on the cell differentiation landscape for a realistic two-gene regulatory network, consisting of self-promoting and mutually inhibiting genes. Our work provides the first theoretical basis of the transdifferentiation process in the presence of delays, where one differentiated cell type can transition to another directly without passing through the undifferentiated state. Additionally, the interplay of time-delayed feedback and a time dependent chemical drive leads to long-lived oscillatory states in appropriate parameter regimes. This work emphasizes the important role played by time-delayed feedback loops in gene regulatory circuits and provides a framework for the characterization of epigenetic landscapes.

## I. INTRODUCTION

The rise of transcription factor mediated induced pluripotency [1–6] has given rise to increased efforts in order to understand the epigenetic landscape that underpins the process of cellular differentiation. Yamanaka and co-authors have shown that four transcriptions factors (TF), Sox2, Oct4, Klf4 and c-Myc are enough to reprogram fully differentiated cells to the pluripotent state, in mouse fibroblast cells [1], as well as in human fibroblasts [2]. A different program, using a combination of the TFs Sox2, Oct4, NANOG, and LIN28 has also been shown to induce pluripotency in fully differentiated human cells [3–5].

The process of cellular differentiation has traditionally been thought of in terms of Waddington’s epigenetic landscape [7], using the metaphor of a ball rolling down a hill, with the final valley representing the differentiated states, the top of the hill representing the pluripotent state, and the valley chosen by a cell from among similar ones as cell fate decisions. The discovery of the reprogramming pathway has led to an interest in mathematically modeling the epigenetic landscape though the underlying Gene Regulatory Networks (GRN). Initial studies using a toy single gene regulatory network studied the properties of cellular differentiation using a self-activating gene [8–12]. Later studies have expanded this work to model more realistic two gene networks, that are self-activating and mutually inhibiting [11, 13–17]. These models have shown that the cell can follow different pathways during the differentiation and reverse programming pathways, and provided a more nuanced understanding of the epigenetic landscape [11, 13–16].

An experimental attempt to elucidate the reverse programming pathway was made by Nagy and Nagy [18], in which they attempted to clarify the pathway of a cell undergoing reverse differentiation through the use of a controlled time-dependent chemical drive. Somatic fibroblast cells derived from mouse iPSC have the four pluripotency factors (Sox2, Oct4, Klf4 and c-Myc) incorporated in a latent state. These were activated through a single chemical factor, doxycycline, which was supplied for different durations. They observed that when the doxycycline input was provided for a time of less than *∼*7 days (*d*_*P*__*NR*_), the cell stayed in the fully differentiated somatic state. On the other hand, on providing the doxycycline input for a period greater than *∼*14 days (*d*_*CP*__*S*_), the cell made a transition to the induced pluripotent state. For a chemical drive between *d*_*P* *NR*_ and *d*_*CP*__*S*_, the system was neither in the initial state, nor did it reach the final pluripotent state, and was stuck in a new uncharacterized state, referred to as the *“Area 51”* state. This provided concrete evidence that the path followed by the cell on the reverse differentiation pathway is not necessarily the same as that during the forward differentiation process.

In a previous work [12], we argued that modeling of the underlying gene regulatory network requires careful consideration of the time delays associated with the feedback regulation of the transcription factors. The reverse differentiation process can take place over a duration of weeks, and is accompanied by a host of changes in the epigenetic markers that characterize the state of the cell, such as changes in histone protein expression levels [19, 20], as well as changes in the chromatin compaction characteristics [21–29]. The timescales associated with these physical changes would impact the feedback regulation, and could have a critical impact in assessing the differentiation pathways. We demonstrated a proof-of-concept of this time-delayed feedback regulation in the context of a single-gene time evolution, where we showed that a competition between the delay timescale and the timescale of the chemical drive can set up long-lived oscillations in the concentration of the transcription factor and interpreted this as a possible explanation of the cells caught in limbo during the differentiation process.

In the current work, we present an analysis of the interplay of time-delayed feedback and a time-dependent chemical drive on a more realistic two-gene regulatory network. The underlying motif of the GRN consists of two self-activating and mutually inhibiting transcription factors, which is an extremely common motif observed in biological GRNs [11, 13–17]. Examples include the the differentiation pathway of the Common Myeloic Progenitor (CMP) cell which differentiates into the Myeloid and Erythroid cell fates depending on the over-expression of the TF PU.1 or GATA1 respectively. The myeloid and Erythroid cell lines differentiate further to produce the different cell fates in the myeloid lineage [30–34]. While a full analysis of the reverse differentiation pathway involves multiple genes mediated by multiple TFs, we restrict our analysis to this smaller subset of two-gene mediated regulatory networks. We show that starting from a differentiated state, it is possible to follow the reverse pathway to reach the undifferentiated state. Depending on the interplay between the chemical drive and the delayed feedback, we observe long-lived oscillatory states in this two-TF GRN, which might provide an explanation of the uncharacterized states observed in reverse differentiation experiments [18]. We also show that this same framework can provide a theoretical basis for the phenomenon of transdifferentiation, where one differentiated cell can switch to a different differentiated cell fate without passing through the undifferentiated or pluripotent state. To the best of our knowledge, this is the first study that provides a theoretical underpinning to the transdifferentiation process, which is well known in experimental contexts [35–42].

In subsequent sections, we first define the differential equations that govern the time evolution of the TFs in the presence of a time-dependent chemical drive and time-delayed feedback. We then present results in different parameter regimes, which show oscillatory states and transdifferentiation events. Further, we characterize the phase space of the model in terms of the drive time and the delay timescale as well as characterize the oscillations observed in greater detail. Finally, we discuss the relevance of the current model in the context of generating a more detailed picture of epigenetic landscapes.

## II. MODEL

Two-gene networks have been studied in the literature as a model system to investigate the properties of epigenetic landscapes. The most common motif in a two gene network is where the transcription factor corresponding to each gene up-regulates its own production and downregulates the production of the transcription factor associated with the other gene (Fig. 1a). It is possible to write down differential equations governing the time evolutions of the concentrations of the two transcription factors,

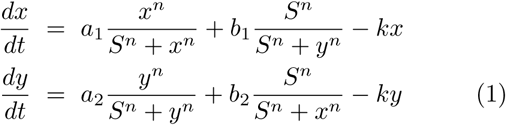

where *x* and *y* denote the concentrations of the two transcription factors [11, 13]. The feedback terms are modeled as hill functions [11], with exponent *n* controlling the steepness of the switch curve, while the parameter *S* controls the concentration at which the half-maximal point is reached. The first term corresponds to a positive feedback mechanism, with each TF up-regulating its own production, while the second term represents the mutual negative feedback between the two components. Finally there is a decay term with the strength of the degradation process controlled by the decay parameter *k.* This model has been studied extensively [11, 13, 16] as a representation of the cell differentiation process and has yielded insights into the epigenetic landscape as a cell differentiates.

**FIG. 1:**
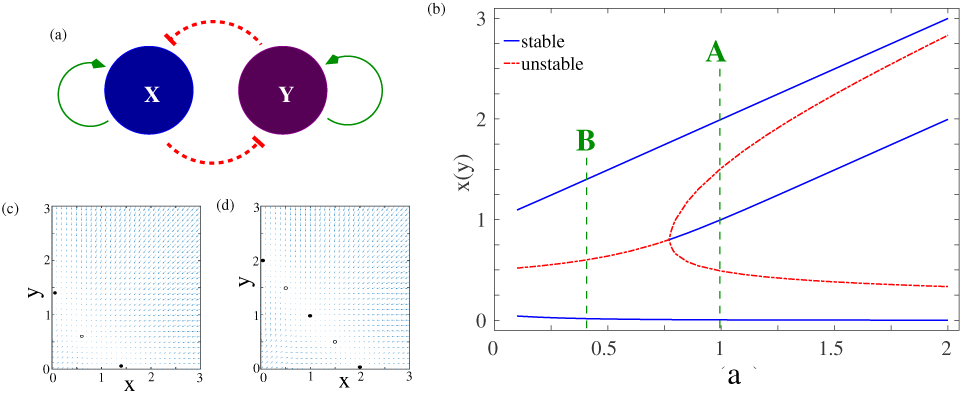
(a) A schematic of gene regulatory network. The two transcription factors X and Y are self-activating, and mutually inhibiting; (b) The bifurcation diagram of the nondelayed system of equations Eq. 1. The dashed green lines correspond to the two values of the feedback parameter *a* that were analysed in subsequent section. The other parameters were fixed at *a*0 = 0.5, *b* = 1.0, *k* = 1.0, *n* = 4, and *S* = 0.5. The velocity vector plots in the (*x, y*) plane for these two points are shown in (c) Regime B and (d) Regime A.

In order to model the reverse differentiation process, we incorporate a time dependent chemical drive (along the lines of our earlier work [12]) as in experiments [18]. The feedback regulation depend on the concentrations of the TFs at some previous times, to account for the finite timescales of chromosome reorganisation and other epigenetic state markers. We model these finite time processes through the incorporation of a time-delayed feedback. The degradation process is still assumed to be dependent on the instantaneous concentration. The time evolution equations incorporating a simultaneous chemical drive and delayed feedback is given by,

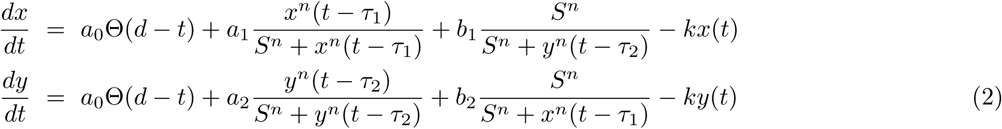

The first term represents the time dependent drive, which is modeled by the Heaviside theta function, with the parameter *d* setting the duration for which the input is provided. Feedback processes mediated by the TF *x* are characterised by a delay timescale *τ*_1_, while those mediated by the TF *y* are characterised by a delay timescale *τ*_2_. The parameters *a*_1*/*2_ and *b*_1*/*2_ controls the strength of the positive and negative feedback for the two transcription factors. In order to reduce the dimensionality of the parameter space, we shall assume them to be the same throughout the remainder of this work, *i.e. a*_1_ = *a*_2_ = *a* and *b*_1_ = *b*_2_ = *b*. The decay constant is assumed to be the same for both TFs for similar reasons.

In the absence of a time-dependent drive and delayed feedback (Eq. 1), the behaviour of the system can be understood in terms of a bifurcation analysis of the phase space. The bifurcation diagram for this system is shown in Fig. 1(b). The phase space analysis shows that this two gene network is a multi-stable system, with a region of bistability for *a < a*_*c*_ *∼* 0.77, and a region of multistability for *> a*_*c*_. For *a < a*_*c*_, the system has two stable steady states (solid blue lines), separated by a central unstable state (dashed red line). The phase space velocities in this region are shown in Fig. 1(c), which clearly shows the two stable attractors separated by the central unstable state. For *a < a*_*c*_, the system undergoes a pitchfork bifurcation where central unstable state become stable and two new unstable states appear symmetrically on either side of the central steady state. The phase space velocity for this region is shown in Fig. 1(d). The central steady state concentrations of the two TFs are equal *i.e.*, *x* = *y*, which in our model corresponds to the undifferentiated (or progenitor) state. The two terminal attractors, one with *x* = *u, y ≈* 0 and another with *x ≈* 0*, y* = *u*, correspond to the two differentiated states. Thus, in each of the differentiated states, one TF completely dominates over the other (*x ≪ y* or *x ≫ y*), with the weaker TF being effectively silenced. These then correspond to two distinct valleys of the epigenetic landscape which represent the two differentiated states. A comprehensive phase space and bifurcation analysis for this non-delayed model has been studied previously [13, 15].

We study the reverse differentiation process, starting from one of the differentiated steady state, and follow the time evolution of the system for different values of the drive duration and delay time. Since the bifurcation diagram shows two distinct dynamical behaviour regimes, *i.e.*, for *a > a*_*c*_ (Regime A), and *a < a*_*c*_ (Regime B), we study the system at a representative point in both these regimes. The parameters chosen for this paper are marked with a dotted line in the bifurcation diagram Fig. 1(b).

## III. RESULTS

### A. Long lived oscillatory states

We first report results for the set of model parameters given by, drive amplitude *a*_0_ = 0.5, the positive and negative feedback amplitudes *a* = 1.0, and *b* = 1.0, decay constant *k* = 1.0, Hill exponent *n* = 4, and *S* = 0.5. We shall refer to this representative point in parameter space as Regime A, as illustrated in the bifurcation diagram Fig. 1(b). The central steady state (undifferentiated state) for this choice of parameters occurs at (*x, y*) = (1, 1) while the two differentiated stable steady states are at (*x, y*) = {(2*, ϵ*) or (ϵ, 2)} with ϵ *≈* 0. The time evolution of the system is shown in Fig. 2 for different values of the drive time *d* for a given value of the delay time *τ*_1_ = *τ*_2_ = *τ* = 500 starting from one of the differentiated states.

**FIG. 2:**
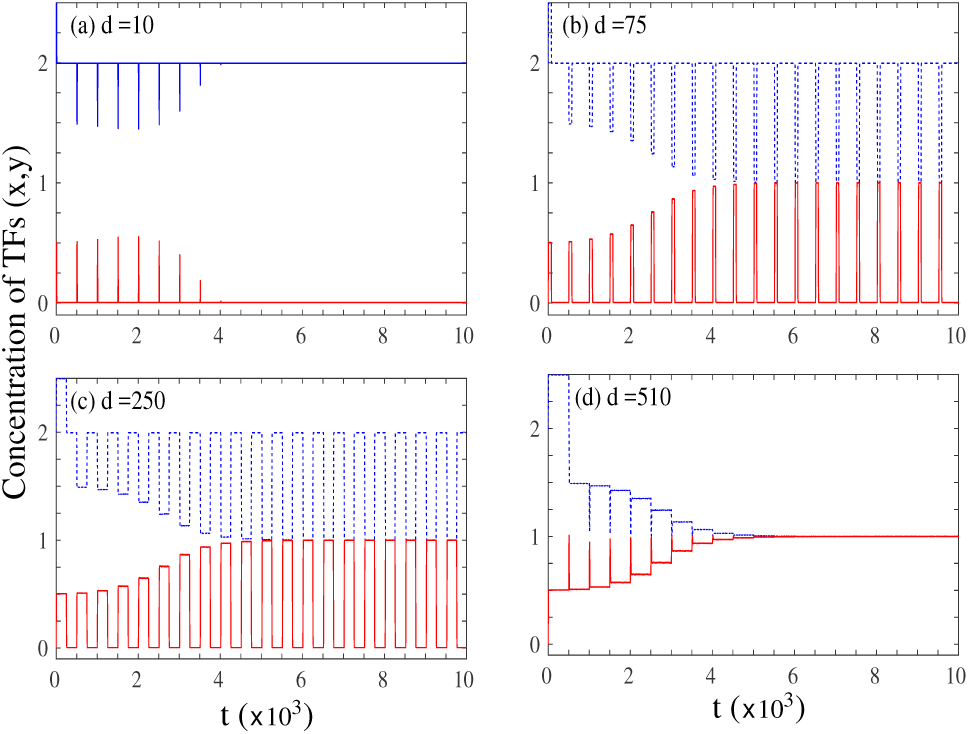
Time series of the concentrations of the two transcription factors (*x/y*) in regime A for four different drive times for *τ* = *τ*1 = *τ*2 = 500. (a) *d* = 10: System stays in initial undifferentiated state; (b) *d* = 75: Sustained oscillations with the system spending more time in the vicinity of one state than the other; (c) *d* = 250: Sustained oscillations with the system spending approximately equal times in the vicinity of the two states; (d) *d* = 510: Successful reprogramming to the undifferentiated state.

Fig 2(a) shows the time evolution of the concentrations of the two TFs when the drive time *d*(= 10)is much less than the delay time *τ* (= 500). In this regime *d ≪ τ*, the concentrations show some initial small fluctuations around the initial conditions, which quickly die out leaving the system in the same state as it initially started with. Thus for small drive times, the system remains in the differentiated state it started out in. As we increase the time for which the drive is provided, the initial small fluctuations grow, until beyond a certain critical value, we observe sustained fluctuations in the concentration of the transcription factors. This is shown for two drive times, *d* = 75 and *d* = 250 in Figs. 2(b) and (c). As can be seen from the figures, for these intermediate drive times, the system oscillates between the initial differentiated state and the central progenitor state (1, 1) for times which are much larger than the characteristic timescales in the system (*d, τ*).

On increasing the drive time even further, the system comes out of this long lived oscillatory state and transitions to central progenitor state beyond a certain critical *d*. This is shown in Fig. 2 for *d* = 510. The system shows small oscillations as it starts from the initial differentiated state, but it quickly settles into undifferentiated state. This would then correspond to a successful completion of the reverse differentiation process. Thus in regime A, when the input is provided for a small time, the system stays in the differentiated state, while if the input if provided for large enough times, it successfully transitions into the progenitor state. In between, for intermediate drive times, the system shows long lived oscillatory states where it is neither in the differentiated nor in the undifferentiated state, but oscillates between the two. The results of our two-dimensional GRN with this interplay of drive and delayed feedback thus closely mirrors the experimental observations of Nagy and Nagy [18], with the identification of the undetermined state in the experiments to the oscillatory state predicted by our analysis.

### B. Transdifferentiation

While the results described for Regime A are robust for *a* > *a*_*c*_ (see bifurcation analysis Fig. 1(b) for details), the system shows a different class of behaviour for *a* < *a*_*c*_. As a representative point in this region of parameter space, we choose *a*_0_ = 1.0*, a* = 0.4, *b* = 1.0, *k* = 1.0, *n* = 5, and *S* = 0.5 and follow time trajectories of the two TFs starting from one of the differentiated states, for different drive and delay times. The drive strength *a*_0_ and the Hill coefficient *n* are changed from the parameter set of Regime A to illustrate the robustness of our results. We shall refer to this representative point in parameter space as Regime B as shown in Fig. 1(b). The (unstable) progenitor state for this choice of parameters corresponds to (*x, y*) = (0.6, 0.6) while the two differentiated states are at (*x*, *y*) = {(1.4, *ϵ*) or (*ϵ*, 1.4)} with *ϵ* ≈ 0.

The time evolution results in regime B is shown in Fig. 3 for different *d* for time delay given by *τ*_1_ = *τ*_2_ = *τ* = 20. When the drive time is small, *d* = 20, the system shows early time transient fluctuations, before relaxing back to the initial state. This is shown in Fig. 3(a), which is similar to the behaviour observed in regime A for small drive durations, where the system does not undergo reverse differentiation, but continues to remain in its initial differentiated state.

**FIG. 3:**
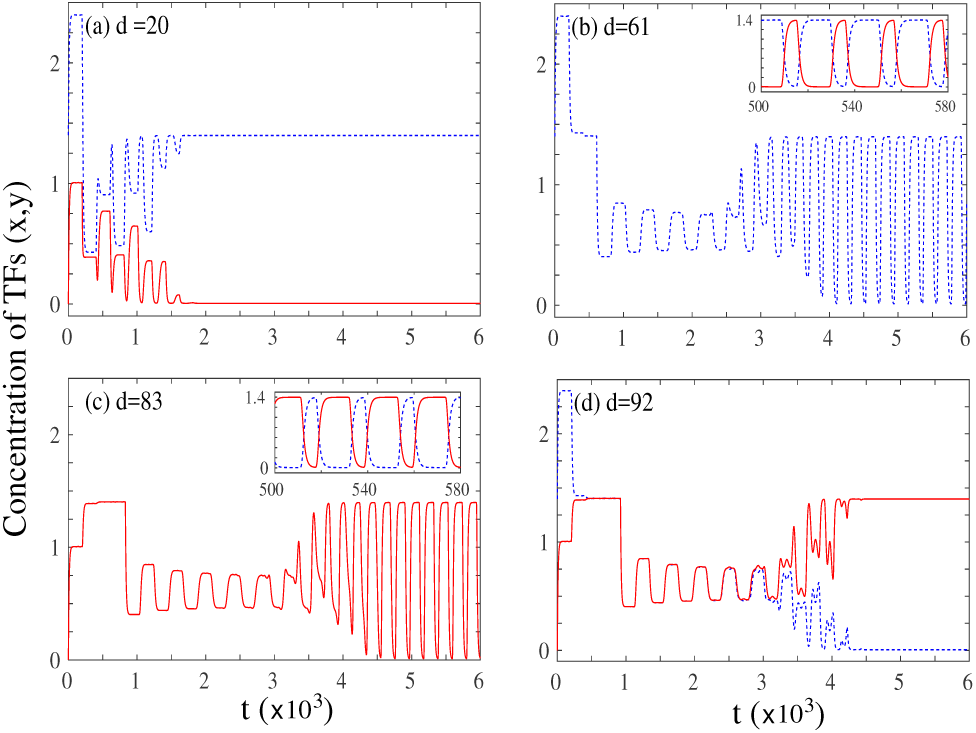
Time series of the concentrations of the two transcription factors (*x/y*) in regime B for four different drive times for *τ* = *τ*1 = *τ*2 = 20. (a) *d* = 20: System stays in initial undifferentiated state; (b) *d* = 51: Sustained oscillations in concentration of TFs. The main panel shows a single TF (*x*) while the inset shows a magnified view of both the TFs in a narrow window of time; (c) *d* = 83: Sustained oscillations in concentration of TFs. The main panel shows a single TF (*y*) while the inset shows a magnified view of both the TFs in a narrow window of time; (d) *d* = 92: Transdifferentiation to the other differentiated state.

On increasing the drive time, the system shows sustained oscillations, as shown in Fig. 3(b) and (c) for *d* = 61 and *d* = 84 respectively. Crucially in this case, the oscillations are between the two differentiated states and not between the initial differentiated state and the progenitor state, as in regime A described above. The oscillations are again long-lived in comparison to the drive and delay time scales of the system. The insets in panels (b) and (c) of Fig. 3 show a magnified view of the oscillations for both the transcription factors.

Finally Fig. 3(d) shows the time evolution of the system for *d* = 82. In this case, after initial transient oscillations, the system stabilizes to a “flipped” state *i.e.* starting from one of the differentiated states (1.4, 0), the system transitions to the other differentiated state (0, 1.4) without reaching the central progenitor state. This dynamics is reminiscent of the phenomenon of transdifferentiation, which has been observed experimentally where cells can directly transition from one differentiated state to another [35–42].

The time evolution observed in regime B is qualitatively different from the behaviour obtained in regime A. In regime B, the system never reaches the central progenitor state for any choice of the drive and delay times. Similarly, the transdifferentiated state is never observed for regime A. Further the oscillatory state is distinct in the two cases, with oscillations between the progenitor and the differentiated state in regime A (half oscillations) and between the two differentiated states in regime B (full oscillations).

To the best of our knowledge this is the first theoretical model that helps elucidate how delayed feedback processes can give rise to transdifferentiation in gene regulatory networks.

### C. Phase diagrams

The different dynamical behaviours in Regimes A and B, characterised by time evolution plots can conveniently be represented as phase diagrams in the drive vs. delay time (*d vs. τ*) plane as shown in Fig. 4. Both situations, where the delay timescale for the two TFs are same *i.e.* (*τ*_1_ = *τ*_2_ = *τ*) and different (*τ*_1_ ≠ *τ*_2_) are considered. The long time behaviour can be classified into four phases (*i*) the system stays in the initial differentiated state (*I*); (*ii*) the system reaches an oscillatory state (*IIA* or *IIB*, depending on the nature of oscillations); (*iii*) the system undergoes reverse differentiation to reach the pluripotent state (*III*); and (*iv*) the system reaches a transdifferentiated state (*IV*).

**FIG. 4:**
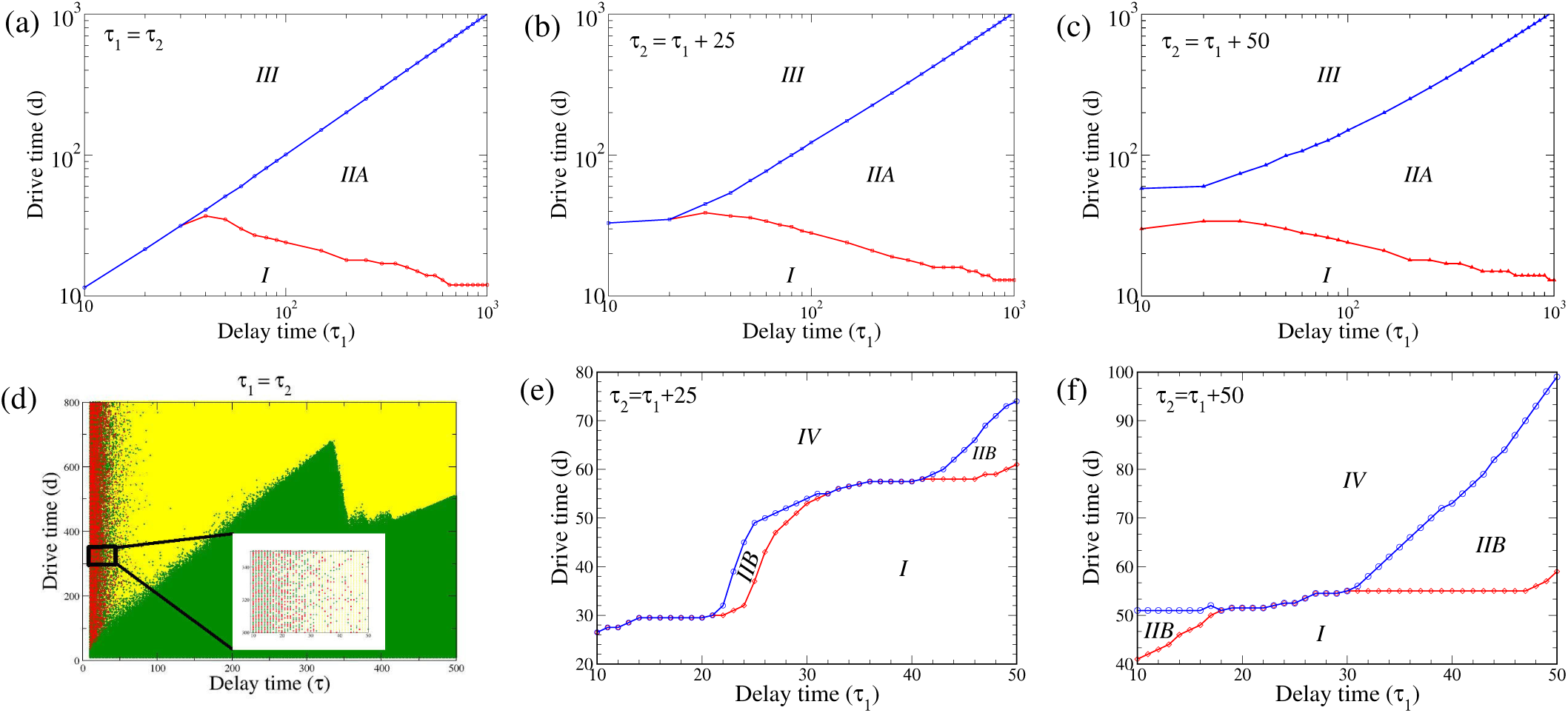
Phase space in the drive time (*d*) vs. the delay time (*τ*1) plane for (a) Regime A: *τ*2 = *τ*1; (b) Regime A: *τ*2 = *τ*1 + 25; (c) Regime A: *τ*2 = *τ*1 + 50; (d) Regime B: *τ*2 = *τ*1. The inset shows a magnified view of a small region of this phase space to illustrate the chaotic behaviour in this regime; (e) Regime B: *τ*2 = *τ*1 + 25; (f) Regime B: *τ*2 = *τ*1 + 50. All other parameters for regime A and regime B are as defined in the text.

Fig. 4(a), shows the phase behaviour of the system in regime A for *τ*_1_ = *τ*_2_ = *τ*. In the region with small delay and drive times *τ < τ*_*cr*_, and *d < d*_*cr*_, the system stays in the initial differentiated state (*I*). For small delay times *τ < τ*_*cr*_, but beyond a critical drive time *d > d*_*cr*_, the system transitions directly from the differentiated state (*I*) to the central progenitor state (*III*), with the critical drive duration for the reverse differentiation process increasing linearly with the delay time *i.e.* (*d*_*cr*_ α *τ*). For *τ > τ*_*cr*_ this direct transition from the initial differentiated to the final undifferentiated state is lost. In this region, the system initially passes from the initial state to an oscillatory state (*IIA*) – as demonstrated in Figs. 2(b) and (c). In line with the experimental results [18], we denote this minimal drive time beyond which the initial state is no longer the stable solution as *d*_*P* *NR*_ (the *Point-of-No-Return*). By keeping *τ* fixed and increasing *d*, the oscillatory state is observed to be stable till a second critical value of the drive time *d*_*CP* *S*_ (*Commitment-to-Pluripotent-State*). Beyond this value of drive time *i.e. d > d*_*CP* *S*_ the undifferentiated state is stable. The threshold for the point-of-no-return *d*_*P* *NR*_ decreases with increasing time delay, implying that the oscillatory state sets in for lower values of the drive time for higher *τ* values. On the other hand, the threshold for crossing to the pluripotent state increases linearly with delay times (*d*_*CP* *S*_ α *τ*), such that the crossover to the central state requires a chemical drive for longer durations as *τ* is increased. Thus with increasing delay timescales, the regime of stability for sustained oscillations continues to increase widening the gap between the point of no return and commitment to pluripotency.

In Fig. 4(b) we show the phase behaviour in regime A for the situation where the delay times are unequal *i.e. τ*_1_ *τ*_2_ with a constant difference between the two delay times, *i.e.* Δ*τ* = *τ*_2_ *- τ*_1_ = 25. Similar to the situation when delay times are equal, the system directly transits from the initial state (*I*) to the final progenitor state (*III*) for small value of *τ*_1_ beyond a critical value of drive time *d*_*cr*_. However unlike equal delay time scenario, the critical drive time at which this transition takes place *d*_*cr*_ remains almost constant as *τ*_1_ is increased. Beyond a threshold *τ*_*cr*_, this direct transition gives way to to a twostep transition, in which stable state is the initial state for *d < d*_*P* *NR*_, the oscillatory state for *d*_*P* *NR*_ *< d < d*_*CP* *S*_, and finally the progenitor state for *d > d*_*CP* *S*_. This is similar to the equal delay time scenario described earlier. Further, the boundaries *d*_*P* *NR*_ and *d*_*CP* *S*_ behave as before, with *d*_*P* *NR*_ decreasing with increasing *τ*_1_ and *d*_*CP* *S*_ increasing linearly with *τ*_1_. The critical value of *τ*_1_ at which this two-step transition take place is lower in this Δ*τ* = 25 case than in the previous Δ*τ* = 0 one. Finally in Fig. 4(c) we show the phase behaviour for the case of Δ*τ* = 50. Interestingly, in this case, there is no value of the time delay *τ*_1_ for which the system makes a direct transition from the differentiated (*I*) to the undifferentiated state (*III*). In this regime upon increasing the drive time *d > d*_*P* *NR*_, the system necessarily passes through a region where the oscillatory state is stable before it can make the transition to the central progenitor state beyond the *d*_*CP* *S*_ drive time. The width of oscillatory region remains constant with increasing *τ*_1_ (unlike the Δ*τ* = 0 and 25 cases) for small values of *τ*_1_ and widens for large values with *d*_*P* *NR*_ and *d*_*CP* *S*_ diverging away from each other.

We now turn to the analysis of the phase behaviour in regime B, *i.e.* transdifferentiation regime. This is shown for *τ* = *τ*_1_ = *τ*_2_ in Fig. 4(d). For small values of the drive time, the system remains in the initial undifferentiated state (*I*). On increasing the drive time, for small values of the delay time *τ* (≲ 50), the final long-term behaviour of the system shows extreme sensitivity to parameter values. The inset of Fig. 4(d) shows a magnified view of the phase map for *τ <* 50 in the region 300 *< d <* 350. As can be seen from the inset, the final state shows signatures of chaotic behaviour, where the long time state is either the initial state (*I* green dots) or oscillatory state (*IIB* yellow dots), or transdifferentiated state (*III* red dots) for slight variations in drive time *d*. The long time behaviour of the system was obtained by time marching the set of delay differential equations (Eq. 2) for a time *t* much greater than the delay and drive timescales of the system (*t* = 10^4^ ≫ {*d, τ}*). While it is possible that the final state obtained from our simulations is not the true steady state, biologically relevant timescales would amount to the states observed after times which are of comparable magnitude to the experimental timescales, such as the time *d* for which the chemical drive is provided to the system [18]. In these parameter regimes, as shown by the phase map, this deterministic system shows chaotic behavior. For larger delay times (*τ ≳*50), the transdifferentiated state is no longer stable (in the range of drive times investigated) and the system transits from the initial state (*I*) to an oscillatory state (*IIB*) beyond a certain threshold drive time *d*_*cr*_. The phase boundary demarcating these two regions shows a non-monotonic behaviour (saw-tooth-like pattern), increasing linearly until *τ* ∼ 340, beyond which it drops sharply before continuing to rise with a different slope.

Interestingly, this apparent randomness is lost when the the two delay timescales are no longer equal, Δ*τ* = *τ*_1_ *- τ*_2_ ≠ 0. Fig. 4(e) shows the phase behaviour in Regime B for Δ*τ* = 25. For small values of *τ*_1_ (*<* 20), the system transitions from the initial state to the transdifferentiated state beyond a critical drive time *d*_*cr*_. For 20 ≲ *τ*_1_ ≲ 30, there exists range of values of *d* which show sustained oscillations. Below this range, the system stays in the initial state, while for larger drive times, the transdifferentiated state is stable. For 30 ≲ *τ*_1_ ≲ 40, there is no oscillatory state, and the system again transitions directly from the initial state to the transdifferentiated state. Above this delay timescale range *τ*_1_ *>* 40, the oscillatory state reappears in the phase map, and the region of stability of the oscillatory state increases with increasing *τ*_1_. While the boundaries between the different phases are sharply defined for regime B in this Δ*τ* ≠ 0 condition, the signature of the chaotic behaviour remains in the appearance and disappearance of the stable oscillatory solution for different drive times.

Finally, Fig. 4(f) show the results for regime B with Δ*τ* = 50. In this case, for low values of *τ*_1_ (≲ 18), the oscillatory state is stable for a region of drive times. In an intermediate regime 18 ≲ *τ*_1_ ≲ 30, there is no stable oscillatory state, and the system transitions from the initial state to the transdifferentiated state beyond a threshold drive time. The oscillatory state reappears for *τ*_1_ ≳30, and as in the Δ*τ* = 25 case, the region of stability of the oscillatory solution increases with increasing *τ*_1_.

The above analysis of the phase behaviour of the model in regime A and B shows that the mutual interplay of drive and delay timescales can give rise to extremely rich landscapes, with non-trivial dependence of the final steady state on the time for which the input is provided, as well as the delay timescales associated with both the TFs. An interesting question to ask is how these exotic dynamical states are realisable in a biological context.

### D. Characterizing the oscillations

We now turn to characterizing the long-lived oscillatory state observed in regimes A and B. In Fig. 5 we show the trajectory of the system in the oscillatory state plotted in the *x*(*t*) vs. *x*(*t* + *τ*) plane. While a stable steady state corresponds to the trajectories evolving to a single point in this plane, steady oscillations evolve to a limit cycle.

**FIG. 5:**
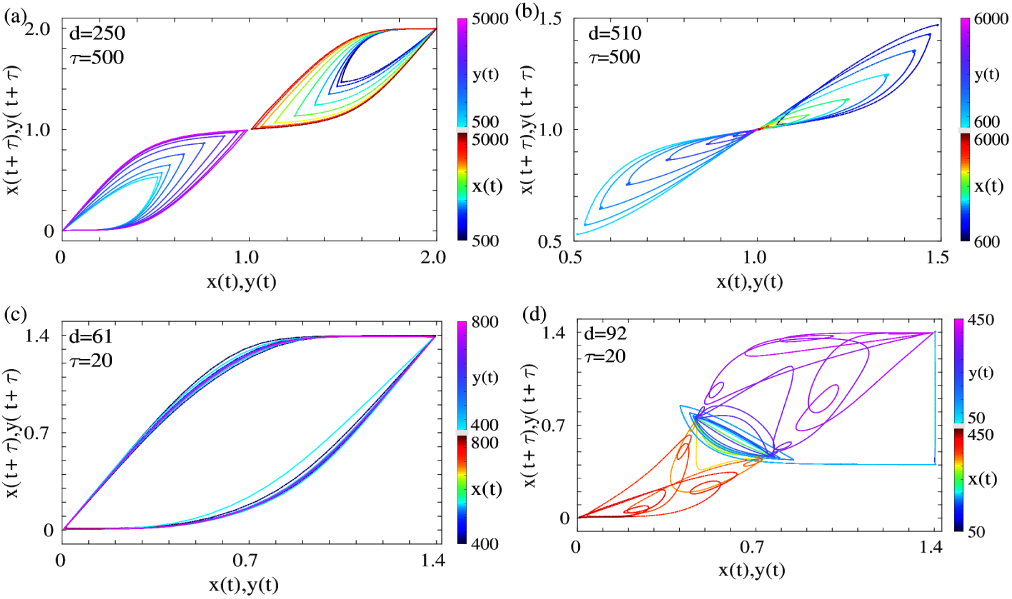
Trajectory plots in the *x/y*(*t*) vs. *x/y*(*t* + *τ*) plane for (a) Oscillatory state in regime A (state *IIA*). The oscillations are between the differentiated states and the central undifferentiated state, which are represented as half-cycles in this phase plane; (b) Reprogramming to the undifferentiated state in regime A (state *III*). Note the decaying transient oscillations in both *x* and *y*, which converge to the central steady state; (c) Oscillatory state in regime B (state *IIB*). The oscillations are between the two differentiated states in this regime, which is represented by single limit cycle in this phase plane; (d) Transdifferentiated state in regime B (state *IV*). Note the unstable oscillations around the central state preceding transdifferentiation. In all four cases, the trajectories are coloured by the time values, as shown in the corresponding colorbars.

Fig. 5(a) shows the trajectory of the system in this plane in regime A for the concentrations of both the TFs *x*(*t*) and *y*(*t*). As can be seen clearly from the time evolution of the trajectories, both the concentrations evolve to a steady limit cycle corresponding to the oscillatory region (*IIA*). The oscillations for each TF are between the initial differentiated state and the central undifferentiated state, which is manifested as two distinct limit cycles for the two TFs in the upper right and lower left quadrant. The corresponding trajectories for regime B are shown in Fig. 5(c). In this case, the oscillations are between the two undifferentiated states (*IIB*), and hence the limit cycles for the two TFs completely overlap, unlike the limit cycle in regime A. In contrast, we show the evolution of the trajectories in this plane for the reprogramming in regime A (Fig. 5(b)) and transdifferentiation in regime B (Fig. 5(d)). For successful reprogramming, initial transient oscillations between the initial state and the central state decay to the central stable state for both the TFs. For transdifferentiation, the concentrations of the two TFs perform small oscillations around the undifferentiated state for a short period of time (blue limit cycle), beyond which the limit cycle becomes unstable and the TFs reach the other differentiated state. The oscillations enable the trajectories to mix, ensuring that the history of the initial state is lost, which triggers the switch to the transdifferentiated state.

While the oscillatory state is stable for *d*_*P* *NR*_ *< d < d*_*CP* *S*_, the nature of oscillations changes as *d* is varied. A quantitative estimate of the difference in the oscillatory state can be obtained by comparing the residence time distributions of the two transcription factors for different drive and delay times. We plot the residence time distribution as a function of the concentration for both regime A (Fig. 6(a)) and regime B (Fig. 6(b)). As can be seen clearly in Fig. 6(a), on increasing drive times, the peak in the probability distribution shifts from a single peak at *x* = 2 for low values of *d* to a bimodal distribution at higher drive times (*d* = 250), and back again to an unimodal distribution, but centered around the undifferentiated state at *x* = 1 at even higher drive times, *d* = 510. In order to estimate whether there is a systemic behaviour across a range of delay timescales *τ*, we plot the probability of finding the TF1 at the vicinity of the undifferentiated state, *x* = 2, against the nondimensionalised drive time *d/τ* for a range of *τ* values. Remarkably, this probability appears to have a simple scaling behaviour dictated by a single parameter *d/τ*, *viz. p*(*x* = 2) = 1 *-* (*d/τ*). While the oscillations in regime B can be characterized by a similar method as shown in Fig. 6(b), the chaotic nature of the oscillatory and transdifferentiated state implies that there is no simple scaling relation with the drive time in this case, unlike in regime A.

**FIG. 6:**
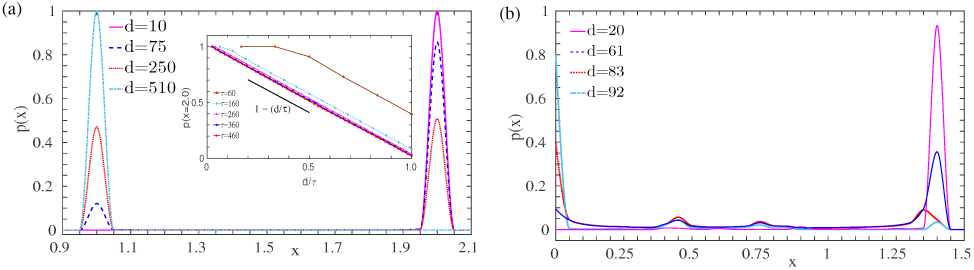
Probability of the concentration of any one transcription factor for different values of the drive time for (a) regime A with *τ* = 500. The inset shows the probability near one differentiated state (*x* = 2) as a function of the ration of the drive to delay times *d/τ*; and (b) regime B with *τ* = 20.

## IV. DISCUSSION

Our work illustrates the richness of epigenetic landscapes in the presence of external drive and delayed feedback. Using a generic model of two self-activating and mutually inhibiting genes, we show that the reprogramming to the common progenitor state requires a delicate balance of the drive and delay timescales, as well as appropriate positive and negative feedback strengths. Apart from the reprogrammed state, the system can find itself in long-lived oscillatory states or in a transdifferentiated state in different parameter regimes. An analysis of the phase diagrams in the delay drive time plane provides a comprehensive picture of the final states of the reverse differentiation process. Additionally, the phase diagrams also provide a signature of chaotic behaviour in appropriate regimes, which have been hypothesized to play an important role in the cell-fate determination process [43]. In the chaotic regime, the model predicts that slight changes in the drive or delay parameters can drive the system from one state to another. The chaotic regime can thus provide a pathway for the control of cell fate determination.

To the best of our knowledge, this work provides one of the first theoretical basis for understanding the phenomenon of transdifferentiation in the presence of delayed feedback. Transdifferentiation was first shown experimentally in mouse fibroblasts in 1987, where the transcription factor MyoD, belonging to the Myogenic Regulatory Factors (MRF) family was shown to induce transdifferentiation of mouse embryonic fibroblasts to myoblasts [35]. The therapeutic potential of transdifferentiation has led to attempts to direct the process towards functional outcomes. The first proof of such a functional transdifferentiation was carried out in mouse liver cells which transitioned to pancreatic-beta-cell-like cells that helped control the effects of hyperglycemia [44]. Transdifferentiation has been shown to be a valid strategy to alter somatic cell fates in humans as well, which was first demonstrated in the transdifferentiation of a human liver cell to human beta cells through the effect of a single gene PDX-1 [45]. The implications of successful transdifferentiation in medical applications has led to a surge of experimental work using a combination of strategies including the forced expression of specific transcription factors and combination of defined factors with microRNAs or small molecules to achieve targeted functional transdifferentiation of one somatic cell type to another [36–42, 46–48]. An understanding of the underlying dynamics that characterizes the transdifferentiation process has however been lacking, including the question of whether cells need to necessarily pass through partially reprogrammed states in order to convert to a different cell type.

The mechanism proposed in this paper articulates one possible pathway for transdifferentiation, in the context of somatic cell types of similar lineages [49]. We show that it is possible to achieve transdifferentiation of one somatic cell type to another without passing through the undifferentiated state. Our analysis also provides an explanation for this process in terms of the underlying bifurcation diagram of the model transdifferentiation arises when the feedback is tuned such that the central progenitor state is unstable. Interestingly, we observe that the transdifferentiation process is preceded by unstable oscillations, as has been seen in a different modeling approach [50]. A different mechanism has been proposed where transdifferentiation proceeds through a distinct stable intermediate state or a series of unstable intermediate states [51]. The incorporation of delays, as in the current work, proposes a new pathway to transdifferentiation, where unstable oscillations around the central state serve as a precursor to the transdifferentiated state. It would be interesting to study the transdifferentiation process in the presence of delayed feedback, in a landscape with multiple lineage branching points in order to characterize the pathway in these more complex situations.

The interplay of delayed feedback and chemical drive also leads to sustained oscillations in the levels of transcription factors. While we predicted the presence of sustained oscillations in the one gene network reported previously [12], this study conclusively highlights the importance of these oscillations for a more realistic two-gene network. Additionally, oscillations in the concentrations of transcription factors show different characteristics, *(a)* oscillations between the differentiated and undifferentiated state in one regime (*IIA*), and *(b)* oscillations between the two differentiated states in a different regime (*IIB*). The two oscillatory states may play different roles in the cellular context. It has been shown experimentally that oscillations in the Notch effector gene *Hes*1 regulates the maintenance of neural progenitor cell types [52, 53]. It has also been reported that Stella shows heterogeneous expression levels and dynamic equilibrium in embryonic stem cells (ESC), which is responsible for the ESC existing in a metastable state where they can shift between Inner Cell Mass (ICM) and epiblast like phenotypes [54]. Oscillations have been reported to be a possible feature of epigenetic landscapes [50, 55]. We show that a possible route to these oscillations is through the incorporation of delayed feedback, and this can potentially explain novel states observed during the reprogramming process [18].

In this paper we consider a *deterministic* model of epigenetic landscapes, and consider the effects of time delays and chemical drive. In cellular contexts, it is also important to re-interpret our results in the presence of stochastic fluctuations, which can play a role in cell fate determination [56–63]. Noise induced transitions between different cell types can play an important role in gaining a comprehensive understanding of the differentiation process. The analysis of stochastic fluctuations in these highly nonlinear and non-autonomous delay differential equations present a significant challenge, and the results from an extensive analysis of the role of noise in these systems shall be presented in a forthcoming publication.

The present work provides a framework for the analysis of reprogramming experiments and underscores the importance in accounting for delayed feedback in building a comprehensive picture of the epigenetic landscape. From the perspective of Waddingtons landscape the inclusion of delayed feedback translates to an asymmetry in the cell fate choices for the forward and reverse reprogramming processes. Models involving a larger number of transcription factors which incorporate the effects of this time delay can help in understanding the reprogramming pathway as well serve as a guide to targeted experiments in manipulating the cell fate determination process.

## Acknowledgments

MKM acknowledges financial support from Ramanujan Fellowship, DST, INDIA (13DST052) and the IIT Bombay Seed Grant (14IRCCSG009). BC acknowledges the Biophysical Sciences Institute, and Institute of Advanced Studies, Durham University for financial support at the time the work was done.

